# Large scale *in vivo* acquisition, segmentation, and 3D reconstruction of cortical vasculature using open-source functional ultrasound imaging platform

**DOI:** 10.1101/2022.03.29.485482

**Authors:** Anoek Strumane, Théo Lambert, Jan Aelterman, Danilo Babin, Wilfried Philips, Gabriel Montaldo, Clément Brunner, Alan Urban

**Affiliations:** Department of Telecommunications and Information Processing - Image Processing and Interpretation, Ghent University-imec, Ghent, Belgium; Neuro-Electronics Research Flanders, Leuven, Belgium; Vlaams Instituut voor Biotechnologie, Leuven, Belgium; Interuniversity micro-electronic center, Leuven, Belgium; Department of Neuroscience, KU Leuven, Leuven, Belgium; Department of Physics and Astronomy, Radiation Physics, Ghent University, Ghent, Belgium

**Keywords:** Brainwide ultrasound imaging, Brain vasculature, Blood vessel segmentation, Stroke

## Abstract

The brain is composed of a dense and ramified vascular network comprising various sizes of arteries, veins, and capillaries. One way to assess the risk of cerebrovascular pathologies is to use computational models to predict the physiological effects of a reduction of blood supply and correlate these responses with observations of brain damage. Therefore, it is crucial to establish a detailed 3D organization of the brain vasculature, which could be used to develop more accurate *in silico* models. For this purpose, we have adapted our open-access functional ultrasound imaging platform previously designed for recording brain-wide activity that is now capable of fast and reproducible acquisition, segmentation, and reconstruction of the cortical vasculature. For the first time, it allows us to digitize the cortical vasculature in awake rodents with a ∼100 µm^3^ spatial resolution. Contrary to most available strategies, our approach can be performed *in vivo* within minutes. Moreover, it is easy to implement since it neither requires exogenous contrast agents nor long post-processing time. Hence, we performed a cortex-wide reconstruction of the vasculature and its quantitative analysis, including i) classification of descending arteries versus ascending veins in more than 1500 vessels/animal, ii) quick estimation of their length. Importantly, we confirmed the relevance of our approach in a model of cortical stroke, which enables quick visualization of the ischemic lesion. This development contributes to extending the capabilities of ultrasound neuroimaging to understand better cerebrovascular pathologies such as stroke, vascular cognitive impairment, and brain tumors and is highly scalable for the clinic.

## Introduction

Functional ultrasound imaging is a cutting-edge neuroimaging modality suited to track subtle hemodynamic changes, as a proxy for neuronal activity (Macé et al., 2011). So far, the extraction of the functional ultrasound signal mostly relies on the integration of multiple cerebral vessels (i.e., penetrating arteries, capillaries, and ascending veins) in user-defined (Urban et al., 2014, 2015b), anatomic-based (Sieu et al., 2015) or single-voxel region of interests (Aydin et al., 2020), based on the local change of red blood cells velocity (Urban et al., 2014, 2015a; Brunner et al., 2022) or threshold activity (Urban et al., 2014, 2015b; Gesnik et al., 2016) and recently on reference mouse (Macé et al., 2018; Brunner et al., 2020, 2021; Sans-Dublanc et al., 2021) and rat brain atlas (Vidal et al., 2021; Brunner et al., 2022). Furthermore, these approaches require a mixing of the blood signal coming from distinct vessels with various functions. Indeed, arteries, arterioles, capillaries, venules, and veins behave and adapt differently to neighboring activity (Shih et al., 2013; Rungta et al., 2021) and general cerebrovascular autoregulation along the cerebrovascular tree, not only in the cortex but also in deeper brain structures (Fantini et al., 2016) - a reality rarely addressed when using ultrasound as neuroimaging modality.

Another great advantage of the used methods is that using Doppler shift in the functional ultrasound images allows for the discrimination of flow directionality without the need for a contrast agent (Macé et al., 2011; Brunner et al., 2022). The extractions of the cerebral vessels from these images would provide more precise information regarding the density of ascending veins and penetrating arteries, the cerebrovascular tree and would support a better characterization of the hemodynamic responses.

This work proposes to extract the cortex-wide vascular tree from a set of brain-wide angiography captured with the functional ultrasound imaging modality. Existing work for complete vessel network extraction relies on dedicated angiography modalities like magnetic resonance angiography (MRA; (Hilbert et al., 2020; Tetteh et al., 2020) or optical coherence tomography (OCT; (Yousefi et al., 2015; Li et al., 2017; Wu et al., 2019). The work by (Cohen et al., 2018) demonstrates that vessel extraction is doable using ultrasound imaging as well, albeit for single vessels that connect to a user-defined point of interest. Other work has demonstrated the extraction of the elliptical intersection of vessels from ultrasound imagery (Guerrero et al., 2007; Wu et al., 2019). A generalized pixel profiling (Babin et al., 2012) and center line extraction procedure (Babin et al., 2018) have proven to robustly detect and extract vessel structures from computed tomography angiography and 3D rotational angiography images.

The ideas behind these last two works (Babin et al., 2012, 2018) are the basis of the vessel extraction of this work. The final method robustly estimates the extent to which individual voxels adhere to the properties of vessels, extracts those that are most likely to be part of vessels and subsequently extracts their centerline to form a 3D skeleton which allows for the analysis of the blood vessel’s morphological characteristics. Such processing is of interest in biomedical image processing, as the morphological characteristics like diameter, tortuosity, and shape of blood vessels are critical for early diagnosis, treatment planning, and evaluation.

## Materials and Methods

### Animals

The rats analysed in this work have been selected from the dataset generated for Brunner et al., 2022. Experimental procedures were approved by the Committee on Animal Care of the Catholic University of Leuven, in accordance with the national guidelines on the use of laboratory animals and the European Union Directive for animal experiments (2010/63/EU). Adult male Sprague-Dawley rats (n=9; Janvier Labs, France) with an initial weight between 200-300g were housed in standard ventilated cages and kept in a 12:12hrs reverse dark/light cycle environment at a temperature of 22°C with *ad libitum* access to food and water.

### Cranial window for brain-wide imaging

The surgical procedure has been previously described in Brunner et al., 2022. Briefly, the cranial window was performed in rats under isoflurane anesthesia (Induction 5%, Surgery 2.0-2.5%, Imaging 1.5% in compressed dry air delivered at 0.6l/min; Iso-Vet, Dechra, Belgium). Xylocaine (0.5%, AstraZeneca, England) was injected subcutaneously into the head skin as pre-operative analgesia. The scalp was shaved and cleaned with iso-betadine before being removed over the entire dorsal skull. The cranial window extended from bregma +4.0 to −7.0mm antero-posterior and ± 6.0mm away from the midline. The skull was carefully removed without damaging the dura. The brain was covered with a low-melting 2% agarose (Sigma-Aldrich, USA) and ultrasound gel to ensure a proper acoustic coupling with the ultrasound probe.

### 2D scans of the brain vasculature with ultrasound imaging

The brain-wide imaging procedure has been previously described in Brunner et al., 2022. Briefly, the functional ultrasound imaging scanner is equipped with custom acquisition and processing software described by Brunner et al., 2021. In short, the scanner is composed of a linear ultrasonic transducer (15MHz, 128 elements, Xtech15, Vermon, France) connected to 128-channel emission-reception electronics (Vantage, Verasonics, USA) that are both controlled by a high-performance computing workstation (fUSI-2, AUTC, Estonia). The transducer was motorized (T-LSM200A, Zaber Technologies Inc., Canada) to allow antero-posterior scanning of the brain. Imaging is performed on an anti-vibration table to minimize external sources of vibration.

The acquisition consisted in cross-section coronal µDoppler image (12.8-mm width, 9-mm depth) composed of 300 compound images acquired at 500Hz. Each compound image is computed by adding nine plane-wave (4.5kHz) with angles from -12° to 12° with a 3° step. The blood signal was extracted from 300 compound images using a single value decomposition filter and removing the 30 first singular vectors (Urban et al., 2015a). The µDoppler image is computed as the mean intensity of the blood signal in these 300 frames that is an estimator of the cerebral blood volume (CBV; Macé et al., 2011, 2013). This sequence enables a temporal resolution of 0.6s, an in-plane resolution of 100×110µm, and an off-plane (thickness of the image) of ∼300µm (Brunner et al., 2021). We performed high-resolution 2D scans of the brain vasculature consisting of 89 coronal planes from bregma (ß) +4.0 to -7.0mm spaced by 0.125mm (**Figure 1**, Step 1).

**Figure 1.**
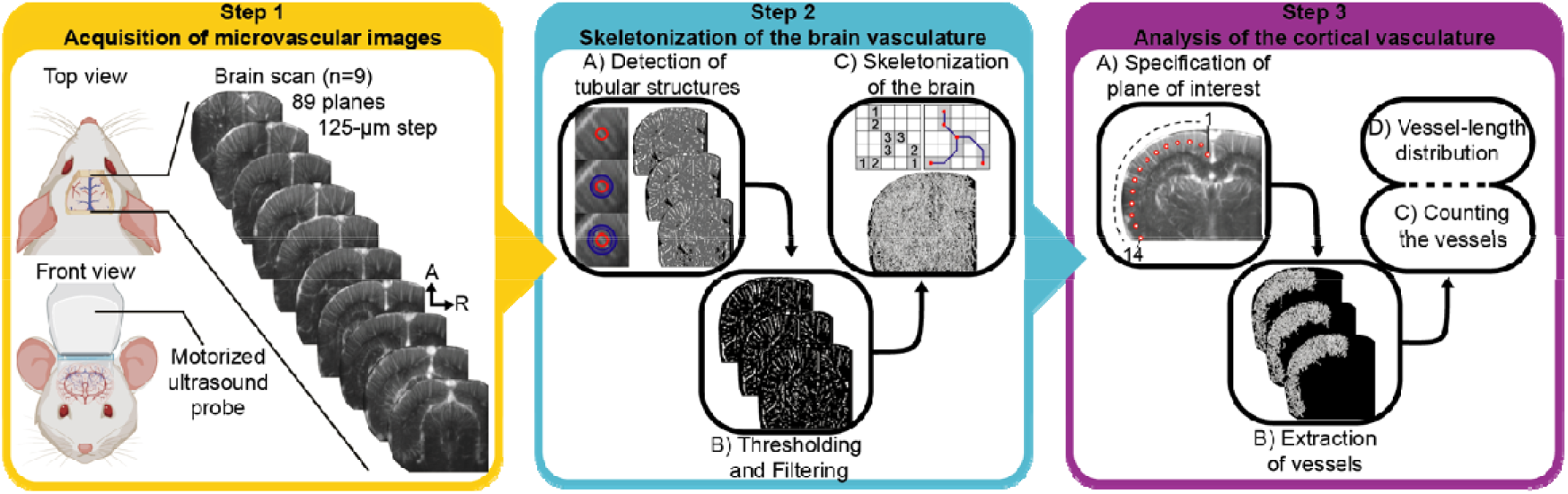
Experimental procedure of whole-brain ultrasound imaging scan of the rat cerebral vasculature (Step 1, yellow), framework for center line extraction and vessel skeletonization (Step 2, blue) and analysis of the cortical vasculature (Step 3, purple). Steps are detailed in the Material and Methods section.

### Stroke

The cortical stroke was induced by the mechanical and permanent occlusion of both the left common carotid artery (CCA) and the distal branch of the left middle cerebral artery (MCA). The post-stroke 2D scan of the rat brain vasculature was performed 70min after the stroke onset. The ultrasound probe was remained in the exact same position between the 2 scans. The procedure to induce the cortical stroke is detailed in Brunner et al., 2022.

### Data processing and analysis

The µDoppler image is acquired and then processed in a cascade of steps: We start with an overview of these steps before explaining in more detail:

1. Step 1: Acquisition of micro-Doppler images of brain vasculature (**Figure 1**, Step 1).
  a. Optional: Color-Doppler filter to separate arteries and veins
2. 2: Vessel extraction and skeletonization (**Figure 1**, Step 2).
  a. Step 2A: Detection of tubular structures using generalized R-profiling: in µDoppler images, vessels appear as a connected series of voxels that are brighter than their surroundings, called ridges. Generalized R-profiling finds and enhances these ridges, making them more suitable for extraction by thresholding.
  b. Step 2B: Filtering of segmented vessels: the enhanced ridges, which indicate vessels, are thresholded to extract foreground voxels. These foreground voxels represent the extracted vessels. As an additional filtering step, isolated foreground voxels, which are most likely to represent noise, are removed.
  c. Step 2C: Skeletonization of the complete brain: the centerlines of the extracted vessels are determined, and a graph type skeleton is built along these lines.
3. Step 3: Analysis of the cortical brain vasculature (**Figure 1**, Step 3).
  a. 3A: Manual specification of plane of interest: a line is drawn in the coronal cross-section µDoppler images of the brain indicating the location of vessels of interest. This line is extended to a plane reaching from posterior to anterior
  b. 3B: Extraction of vessels traversing plane of interest
  c. 3C: Counting of the number of vessels
  d. 3D: Analysis of vessel length distribution

The most complex data processing steps are steps 2A and 2C, which we now explain in greater detail.

### Detection of tubular structures using generalized R-profiling

The goal of “Step 2A: Detection of tubular structures using generalized R-profiling” is to calculate per-voxel measures that express how contrasted a voxel is compared to its multi-scale neighborhood. We achieve this by comparing the voxel gray value to the values found in its neighborhoods (for different sizes). This is based on the *R-profiling* approach described in Babin et al., 2018. The neighborhoods used in this paper are spherical structures with a radius ranging from 1 voxel to 5 voxels, chosen to correspond to the radius of the largest vessels. From each sphere of radius r, the sphere of size r-1 is subtracted, to result in a differential spherical structure. The new voxel value (called the profile measure) is the number of consecutive neighborhoods (for increasing size of neighborhoods) for which the current voxel is brighter than or as bright as its neighborhood. For example, this brightness can be defined as the mean voxel gray value of a neighborhood. The voxel gray value of a central voxel is then compared to the mean gray value of its neighborhood of radius r = 1 and if it is greater than or equal to this mean value, the process is repeated for r=2 and so forth. The largest radius for which this condition is fulfilled (up to the maximum value r=5), becomes the new voxel value (**Figure 1**, Step 2A).

The brightness that is associated with a neighborhood may be chosen using a variety of basic aggregation operators for the neighborhood’s voxel values like mean, maximum, minimum, median, etc. The choice of these operators is made by considering both the properties of the image (such as noise and artifacts) and the type of structures that need to be extracted. It is even possible to define the operator as the average of multiple other operators. The chosen operator is called the profile function.

As the goal of this paper is to extract and analyze vessels, an operator that promotes the ridge-like appearance of vessels is defined. The different basic profile functions that were used are listed below:

1. Maximum: the maximum value,
2. Median (+): the median, taking the highest value if there are two medians,
3. Median (-): the median, taking the lowest value if there are two medians,
4. Minmax average: the mean of the minimum and maximum,
5. Minmax root: the root of the multiplication of the minimum and maximum,
6. Median root: the root of the multiplication of median (+) and median (-).

To evaluate robustness when gathering analytical information, three different operator functions are used on each rat. These three operator functions were respectively an average of the following three combinations: [1,2,3], [1,4,5], and [1,5,6].

Applying these three different profile functions on a µDoppler image of a rat brain results in three 3D images with voxel values ranging from 0 to 5, where the detected vessel-like structures have a higher value than the non-vessels. These images are now ready to be thresholded to result in an extraction of the detected vessel-like structures.

### Skeleton creation

To extract the centerline of the resulting vessels, ordered skeletonization is applied as proposed in Babin et al., 2012. For all foreground voxels, the Euclidian distance to its nearest background voxel is determined. The voxels are then sorted by their respective distances in an ascending order. The voxels are iterated over in this order and a voxel is removed if considered redundant to maintain skeletal connectivity. In other words, a central voxel is redundant in its 27-neighborhood if removing it does not change the number of connected components in that neighborhood. The remaining foreground voxels represent the centerlines of the extracted vessels.

The remaining foreground voxels in the centerline image are subsequently labeled according to their number of foreground neighbors, yielding the following label values (**Figure 1**, Step 2C):

- 0: represents an isolated voxel and is not considered further,
- 1: represents and endpoint of a vessel, and will be represented as a node in the graph-type skeleton,
- 2: represents a part of a vessel, and will be represented as a link in the graph-type skeleton,
- >2: corresponds to a vessel bifurcation. All connected voxels with a degree larger than two represent a single vessel bifurcation. The geometric median of these voxels is used as the location of the bifurcation, thus resulting in a single node in the graph-type skeleton.

The resulting graph consists of vessel endpoints or bifurcations represented by nodes and vessel branches represented by links.

## Results

In this study, we have segmented the rat brain vasculature based on the µDoppler signal captured with ultrasound imaging. We have automatically reconstructed a 3D skeleton of the entire cerebral microvasculature (**Figure 1**). We then extended the analysis to identify and classify individual cortical vessels as penetrating arterioles or ascending venules based on their flow directionality away or toward the ultrasound transducer, respectively (**Figure 2**). Finally, we performed the 3D reconstruction of the cortical microvasculature in a pathological context of cortical stroke (**Figure 3**). The global procedure from brain-wide scan, vessel extraction, and analysis is depicted in **Movie 1**.

**Figure 2.**
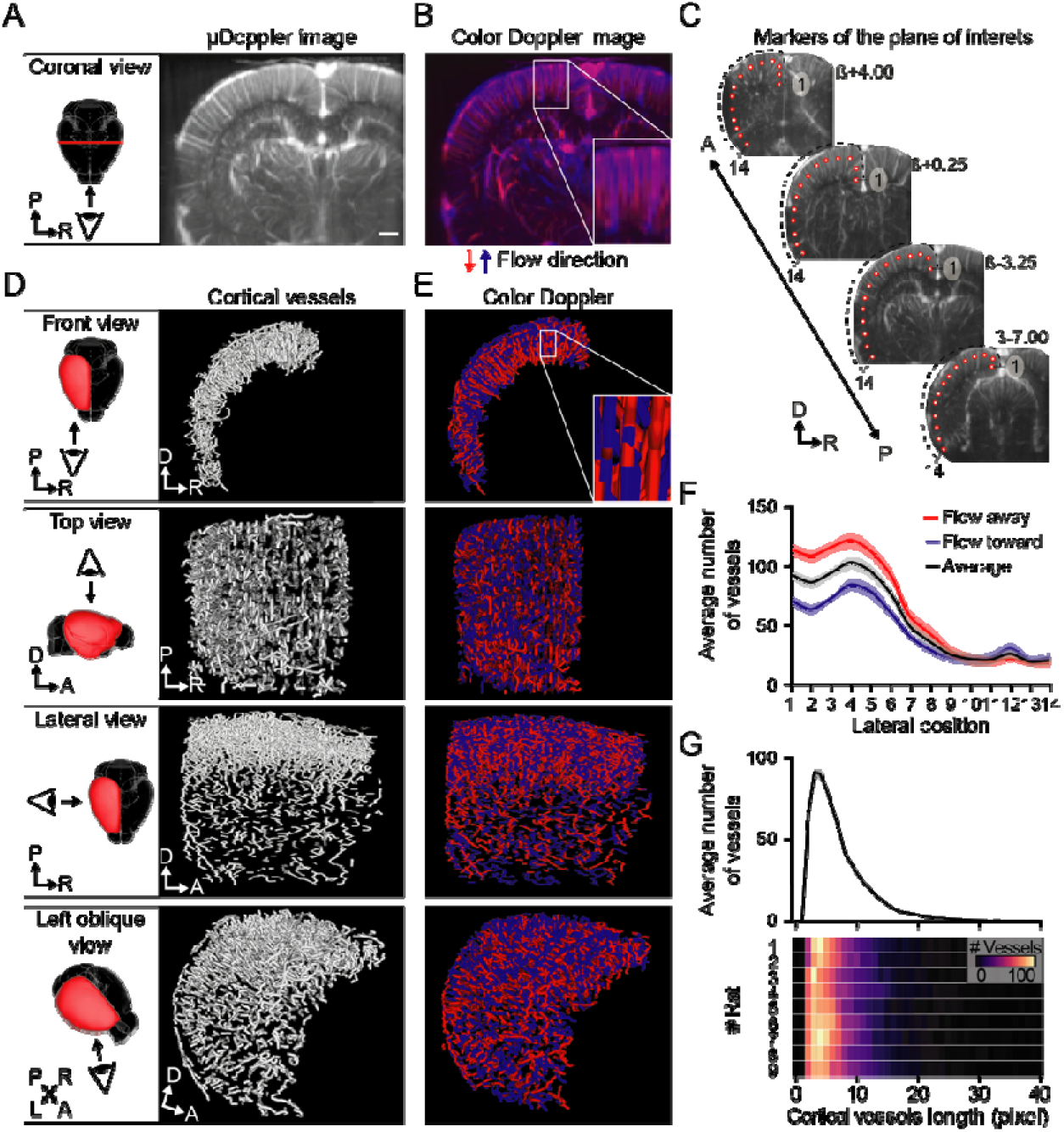
**(A)** Coronal cross-section µDoppler image and **(B)** color-coded image depicting flow directionality toward (blue) or away from (red) the ultrasound probe extracted from the same brain scan. **(C)** Position of markers (1 to 14 from medial to lateral) used to perform the vessel extraction. **(D)** Front, top, lateral, and left-oblique views (top to bottom) of the 3D volume-rendered skeleton of all and **(E)** flow-discriminated cortical vessels. **(F)** Average number of cortical vessels (mean ± sem) and **(G)** average (top; mean ± sem, n=9) and individual distribution (bottom) of cortical vessel length. R, right; L, left; P, posterior; A, anterior; D, dorsal. Scale bar: 1mm.

**Figure 3.**
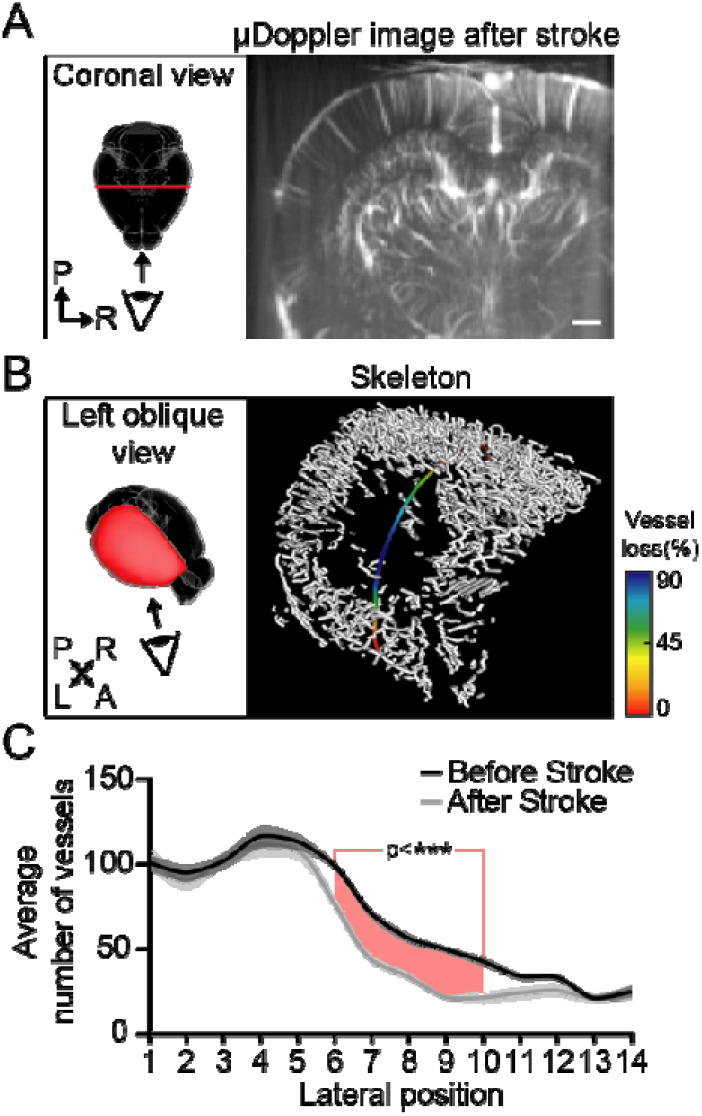
**(A)** Coronal cross-section µDoppler image extracted from the brain scan after stroke. This coronal µDoppler image is at the exact same brain position as in Figure 2A. **(B)** Left-oblique view of the 3D volume-rendered skeleton of cortical vessels after stroke and average drop in vessels detection along the antero-posterior axis (% of vessel loss; color coded line). **(C)** Difference of cortical vessels number along the medio-lateral plane of interest before (black) and after stroke (grey). Positions of the medio-lateral plane of interest depicting significant differences (with a pvalue<0.001***) are highlighted in red. R, right; L, left; P, posterior; A, anterior; D, dorsal. Scale bar: 1mm.

### Ultrasound imaging for skeletonization of the cortical vasculature

Brain-wide imaging was carried out in head-fixed anesthetized rats (n=9) directly after being subjected to cranial window surgery. An angiographic scan composed of a set of 89 μDoppler coronal slices, spaced by 0.125 mm, was performed using a motorized ultrasound transducer (**Figure 1**, Step 1). From this brain-wide scan, we have extracted individual coronal cross-section of µDoppler image based on the relative CBV signal (**Figure 2A** and **Movie 1**). Then, we have skeletonized the brain vasculature following the cascade of image processing steps described in the methods section (**Figure 1**, Step 2). The resulting structure is a 3D skeleton of all the vessels in the brain that were in the field of view of the ultrasound probe. Vessels of interest, specifically the cortical vessels of the left hemisphere, were then extracted by means of a user-defined plane of interest positioned below the brain surface (**Figures 1-3A, 1-3B, 2D** and **Movie 1**). To count the number of vessels in the extracted region of the brain, the plane of interest was divided into 14 sections of equal length in the medio-lateral direction. Per section, the vessels traversing their region of the plane of interest were counted (**Figure 1-3C**). These vessel counts are prone to a slight error, as longer vessels can pass through the plane of interest in more than one of these sections, resulting in a slight overestimation of the total number of vessels. A similar source of slight error is that the point spread function causes some vessels to be detected in multiple planes of interest due to its blurring effect. To determine the distribution of the vessel lengths, the pixel length of each vessel in the extracted skeleton of interest was calculated (**Figure 1-3D**). Because the voxel size in the μDoppler coronal images is anisotropic, this pixel length should not be interpreted as a one-to-one map to the actual vessel’s length, but rather a correlated quantity. The exact measurement of the length remains limited by the anisotropic resolution of the ultrasound imaging voxel technique.

The Doppler measurements allow to generate color-coded images based on the blood flow directionality (**Figure 2B**) as well. This representation of the brain microvasculature allows for the vessel discrimination between penetrating arteries and ascending veins in the cortex. Still, the Doppler measurement is not perfectly accurate and does not result in a 3D velocity vector for the flow, rather yielding a projection of the vector. this nonetheless results in two separate images, one clearly depicting the penetrating arteries, the other the ascending veins. These two images were separately processed in the same manner as the original μDoppler images (i.e., with no separation between veins and arteries). The resulting vein and artery skeletons were overlayed to visualize the combination of both (**Figure 2E**). Due to the aforementioned resolution issues, the same vessels can appear in both images, resulting in the vessel being counted as both a vein and an artery. For this reason, the total number of veins and arteries is slightly overestimated.

From this processing we have counted 892.6 ± 57.5 cortical penetrating arteries (i.e., flow direction away from the ultrasound probe; mean ± sem, n=9) and 641.7 ± 53.1 cortical ascending veins (i.e., flow direction toward the ultrasound probe; mean ± sem, n=9). We have observed an unequal distribution of cortical vessels along the medio-lateral plane of interest ranging from 114.6 ± 5.2 cortical penetrating arteries medially (i.e., position 1) to 18.0 ± 3.9 laterally (i.e., position 14; **Figure 2F**, red plot) and respectively 71.6 ± 4.4 and 23.0 ± 5.1 for ascending veins (**Figure 2F**, blue plot). The medio-lateral variability can be explained by i) the uncertainty of the speed estimator in the estimation of flow directionality (see zoom-in views of color Doppler image in **Figure 2B** and skeleton in **Figure 2E**), and ii) the probe-to-cortical vessel angle potentially affecting the detection of flow directionality (Brunner et al., 2022). However, the number of penetrating and ascending vessels counted along the antero-posterior axis and among animals is highly robust (**Figure 2F**). Moreover, the length of the cortical vessels computed from cortex-wide skeletons follows a beta distribution centered in 3-pixel long vessels (**Figure 2G**, top), representative for all rats processed (**Figure 2G**, bottom).

### Cortex-wide skeletonization for precise detection of ischemic stroke

In a second and pathological application of the center line extraction, we have scanned the rat brain 70min after they have been subjected to concomitant CCA+MCA occlusions and performed the skeletonization of the cortical vasculature as described above. The cortical stroke was confirmed by the direct visualization of the coronal cross-section µDoppler images showing a distinct loss of signal onto the cortex of the left hemisphere when compared with pre-stroke scan (**Figure 3A, Supplementary Figure 1**, and **Movie 3**. See details in (Brunner et al., 2022). In a representative case, the skeletonization of the cortical vasculature after stroke allows the precise detection of the ischemic territory (**Figure 3B** and **Movie 3**) and showing a nearly 90% drop in detected vessels at the center of the ischemic regions along the antero-posterior (**Figure 3B**, color-coded line), but is significantly reduced of ∼50% between lateral position 6 to 10 (grey curve, **Figure 3C** and **Supplementary Figure 2**; pvalue<0.001*** obtained by ordinary two-way ANOVA and uncorrected Fisher LSD with single pooled variance), i.e., where the vascular territory supplied by the distal branch of the MCA is located.

## Discussion

The experimental results show that the µDoppler ultrasound technique yields imagery of sufficient quality to extract vasculature connectivity through the proposed skeletonization method. This enables subsequent statistical analysis of the extracted vasculature. The extraction of these cortical vessels supports a better characterization of the functional ultrasound imaging signal based on vessel length, density, blood speed and thus also offering the measurement of the cerebral blood flow (CBF).

An observed point-of-attention for the vasculature extraction is an overestimation of the number of vessels. This happens due to three reasons: i) vessels traversing multiple segments of the plane of interest, ii) the spatially varying voxel resolution of the ultrasound imaging technique and iii) the uncertainty of the speed estimator (in the case of separation between veins and arteries). The first cause only has effect when the plane of interest is segmented into multiple regions. This issue will not take place if the total number of vessels of the complete plane is being counted, as every vessel will only be counted once. A way to counteract this over-estimation in the different segments is to only count vessels when they are seen traversing the plane for the first time. However, then the ratio of the vessel density throughout the different segments would be less accurate. The other causes may be mitigated through improved HW resolution and/or multi-image fusion to extract full 3D velocity profiles.

At the current stage, the vessel lengths are given in pixels. These measurements cannot be interpreted as exact values because the pixel size is not isotropic. A solution for this would be to take the direction of vessels into account when calculating their length. Knowing the exact length, width and height of the pixels, these lengths may be directly converted to a more meaningful length metric, rather than number of pixels.

Throughout the processing of the images only one step requires user interaction: the specification of the plane of interest. In future application, this step could be done automatically, making the method even more robust when comparing the results of different rats. For example, a brain segmentation technique could be used to find the outline of the brain, which allows the plane of interest to be placed automatically at a fixed distance from this outline. In this way, once the method has been tuned for its specific application, the process can be done automatically without the need of any user interaction.

In terms of resolvable vessel resolution, the data processing procedure produces results at the resolution of the fUS image, in other words: single-voxel-width vessels may be extracted by this procedure. The processing complexity is dominated by step 2A. This is a numerical procedure of linear time complexity in the number of voxels being processed. The implementation developed for the experiments in this paper takes 4-5 minutes on a single-core CPU platform, for a 2-megapixel voxel fUS image. Specific code optimization and implementation on a parallel computing platform (GPU / FPGA) can realistically be expected to bring this processing time down to the second range, opening the door to interactive, real-time vessel imaging and quantification using the fUSI modality.

A direct application of the center line extraction would be to use the skeletonized brain-wide vasculature as a new strategy to improve the filtering, better characterize the fUSI signal and better understand the cerebral hemodynamics and neurovascular coupling. Indeed, one could use the skeleton as a mask to automatically extract hemodynamic parameters from veins and/or arteries such as cerebral blood volume (CBV), red blood cells velocity and even the cerebral blood flow (CBF) in a straightforward and quantitative parameter. Furthermore, it would also allow for live detection of vessels and hemodynamics in preclinical/clinical context. In the meantime, our 3D reconstruction approach combined with the recent improvement vascular ultrasound super-resolution imaging without contrast agents (Bar-Zion et al., 2021), would allow to better separate the dense and overlapping blood vessels; thus, performing a complementary analysis focusing on 2^nd^/3^rd^ order arterioles, venules, and capillaries smaller than the resolution of the functional ultrasound imaging modality, but where neurons mostly modulate blood supply (Rungta et al., 2021).

Moreover, the brain-wide skeleton could help improve the registration by providing a better fit with the reference atlas of the model investigated (Macé et al., 2018; Brunner et al., 2020, 2022; Takahashi et al., 2021) either by the means of vascular landmarks combined with automatic alignment approaches (Nouhoum et al., 2021) or markerless CNN-based deep learning classification (Lambert et al., 2022).

Several strategies are now allowing for longitudinal monitoring of the brain functions of the same animal which in this context would be of high interest when considering pathologies as the brain vasculature and the associated functions have been shown affected (e.g., angiogenesis, vascular rarefaction, hypertension) in various diseases including Alzheimer’s (Meyer et al., 2008; Gutierrez et al., 2016; Zhang et al., 2019; Lowerison et al., 2021; Szu and Obenaus, 2021) and Parkinson’s diseases (Yang et al., 2015; Al-Bachari et al., 2020; Biju et al., 2020), tumors (Gambarota et al., 2008; Guyon et al., 2021), obesity (Dorrance et al., 2014; Pétrault et al., 2019; Gruber et al., 2021) currently addressed at the brain-wide scale with either technology offering reduced spatiotemporal resolution (Pathak et al., 2011; Lin et al., 2013) or *post-mortem* strategies (Hlushchuk et al., 2020; Todorov et al., 2020; Bumgarner and Nelson, 2022). In the stroke context, the cortex-wide skeletonization of vessels could also be used as a follow-up strategy for detecting i) progressive and long-lasting neovascularization of the infarcted tissue (Ergul et al., 2012; Liman and Endres, 2012), and ii) stroke-induced of reverse flow (Li et al., 2010; Ergul et al., 2012) both of crucial interest when considering tissue survival and functional recovery of the insulted tissue.

This paper validates the digitization of rat brain vasculature from ultrasound imagery without the need for contrast agent. It presents an image processing procedure for extraction of vessel structures that was shown to be robust throughout different test subjects while needing little user interaction. The complete process shows promising results with the possibility of acceleration to work in near real-time. Cortex-wide quantification in terms of vessel number and length and discrimination between veins and arteries was successfully shown. The use of such quantification was demonstrated in the case of ischemic stroke, where a clear drop in number of vessels was observed in the expected region.

## Supplementary Information

**Supplementary Figure 1.**
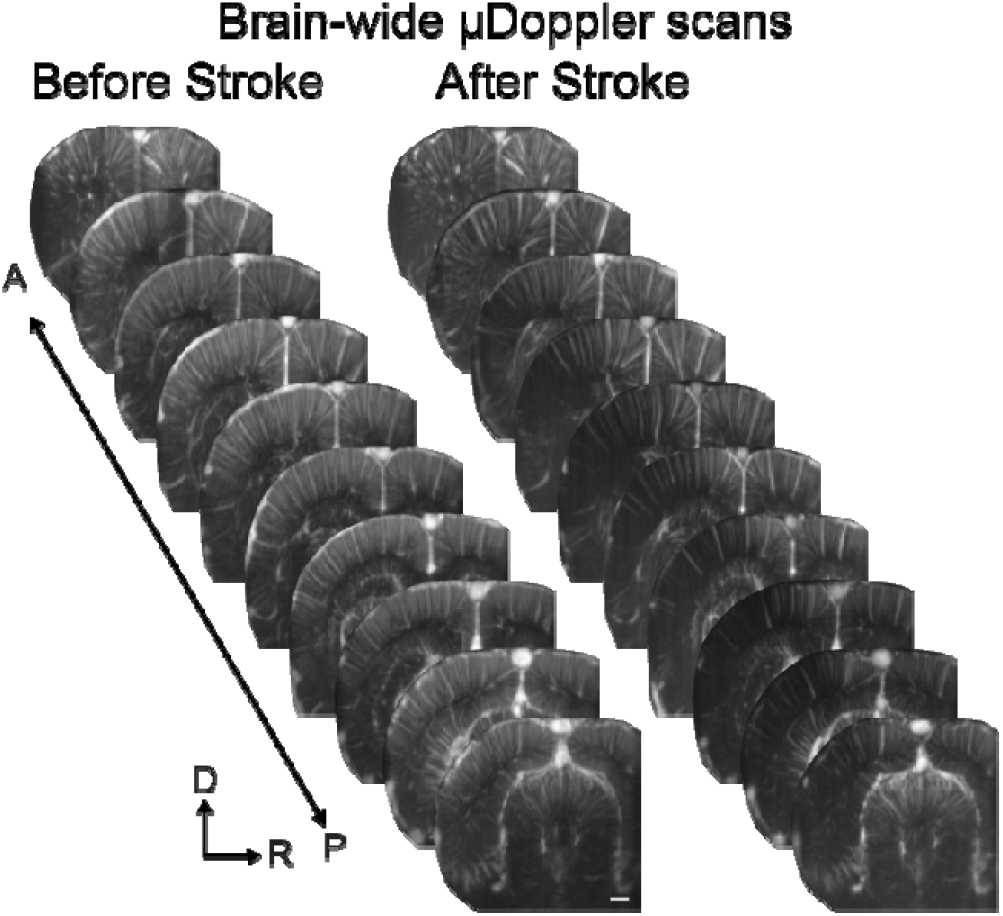
Related to Figure 3A-B. µDoppler scans of the cortical vasculature before and after stroke. Scale bar: 1mm.

**Supplementary Figure 2.**
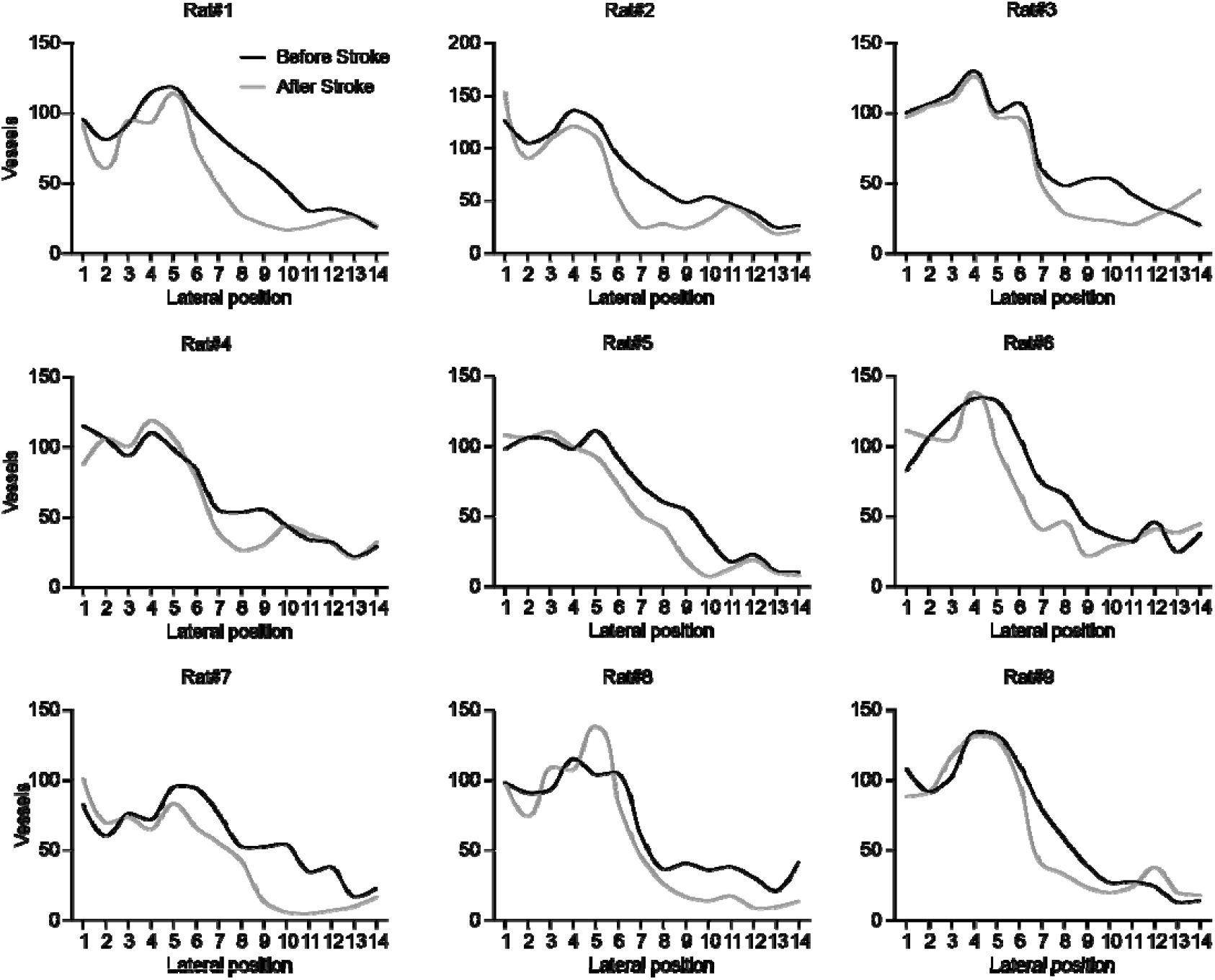
Related to Figure 3C. Number of cortical vessels along the medio-lateral plane of interest before (black) and after stroke (grey) for every single rat used in this work.

**Movie 1**. Experimental pipeline from whole-brain ultrasound imaging scan of the rat cerebral vasculature to vessel skeletonization.

**Movie 2**. 3D volume-rendered skeleton of flow-discriminated cortical arteries (red) and veins (blue).

**Movie 3**. 3D volume-rendered skeleton before and after cortical stroke.

## Data Availability Statement

The dataset and code are available on Zenodo and can be downloaded at https://doi.org/

## Author’s contribution statement

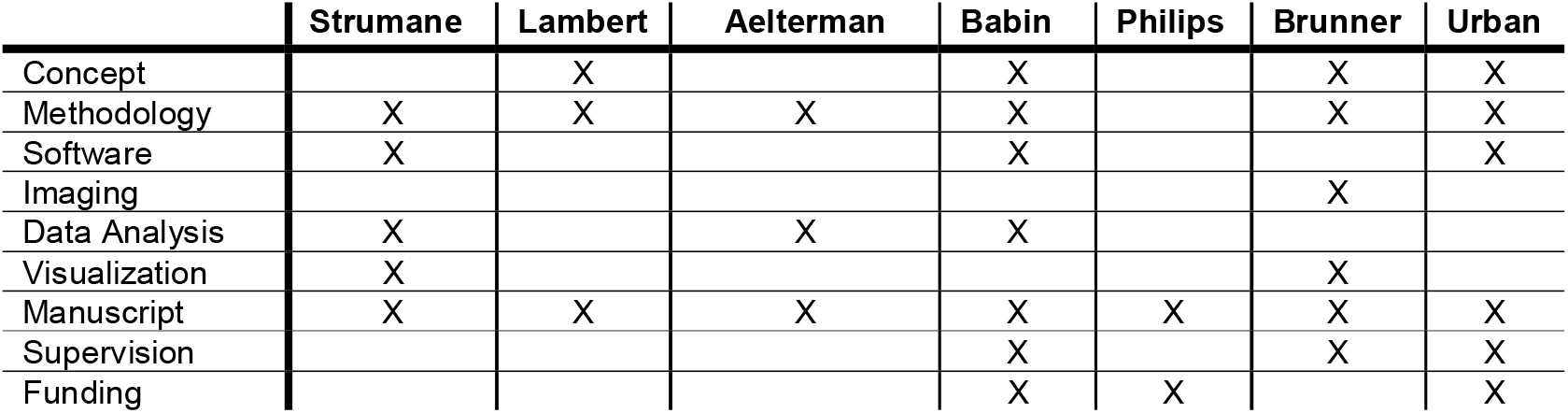

AS: Methodology, Software, Data Analysis, Visualization, Manuscript.

TL: Concept, Methodology, Manuscript.

JA: Methodology, Data Analysis, Manuscript.

DB: Concept, Methodology, Software, Data Analysis, Manuscript, Supervision, Funding.

WP: Manuscript, Funding.

CB: Concept, Methodology, Imaging, Visualization, Manuscript, Supervision.

AU: Concept, Methodology, Software, Manuscript, Supervision, Funding.

## Acknowledgements

We thank all NERF animal caretakers, including I. Eyckmans, F. Ooms, and S. Luijten, for their help with the management of the animals.

